# Analysis of intracellular and intercellular crosstalk from omics data

**DOI:** 10.1101/2023.08.10.552776

**Authors:** Alice Chiodi, Paride Pelucchi, Ettore Mosca

## Abstract

Disease phenotypes can be described as the consequence of interactions among molecular processes that are altered beyond resilience. Here, we address the challenge of assessing the possible alteration of intra- and inter-cellular molecular interactions among gene sets, which are intended to represent processes and or cellular phenotypes. We present an approach, designated as “Ulisse”, which complements the existing methods of enrichment analysis and cell-cell communication analysis. It can be applied to a gene list as well as multiple ranked gene lists, typically derived in the context of omics or multi-omics studies. The approach highlights the presence of alterations in those components that control the interactions between processes or cells. Crosstalk quantification is supported by two null models. Further, the approach provides an additional way of identifying the genes associated with the phenotype. As a proof-of-concept, we applied Ulisse to study the alteration of pathway crosstalks and cell-cell communications in triple negative breast cancer samples, based on single-cell RNA sequencing. In conclusion, our work supports the usefulness of crosstalk analysis as an additional instrument in the “toolkit” of biomedical research for translating complex biological data into actionable insights.

## INTRODUCTION

The understanding of how gene-related molecular alterations translate into pathological phenotypes is a major challenge in life sciences. The experience gained from reductionist approaches like genome-wide association studies – where millions of single nucleotide variations are independently tested for association with a phenotype – strengthen the key role of molecular *interactions*^*1*^. The term “network medicine” indicates the application of network science to study diseases, which are viewed as the consequence of molecular alterations on a complex system of interacting molecular processes^2^.

At cellular level we can classify molecular interactions into two broad categories: intra-cellular and inter-cellular, according to whether they take place *within* a cell or *among* cells. Even if our knowledge of intra- and inter-cellular molecular interactions is incomplete, molecular networks are a crucial tool in biomedical research to translate complex molecular data from omics and multi-omics studies into actionable results^3–5^.

Here, we address the challenge of assessing the possible alteration, with respect to a reference condition, of intra- and inter-cellular molecular interactions among sets of genes, which are intended to represent intra-cellular or cellular phenotypes. There are several differences between our approach, designated as Ulisse, and those already proposed in the field of network medicine (**Supplementary Table 1**). Ulisse can be applied to analyse gene list(s) or ranked gene list(s), two general formats that accommodate any score derived from omics data. We provide a means to screen the alteration of intra-cellular pathway crosstalks and derive a map of the altered communications among pathways, which complements pathway enrichment analysis. The availability of computational tools to study pathway crosstalks is limited, despite their importance in regulatory mechanisms^6,7^, obtaining effective drug combinations in cancer^8^ and investigating complex diseases phenotype^9^. Moreover, Ulisse can be used to reconstruct a cell-cell communication network between cell types/clusters. These two analyses (intra- and inter-cellular) can be combined to obtain integrated pathways of interactions that associate cell-cell communications with intracellular states. Further, we provide a score and a statistical assessment of the altered interactions, based on multiple empirical null models for networks. Lastly, we extract the key genes that take part in the altered interactions.

**Table 1.** Top 5 pathway crosstalks mediated by cancer cell DEGs. The genes reported between parentheses are the DEGs that contribute to the crosstalk. The notation | · | indicates gene set size, while ‖·‖ indicates the sum over all gene weights that contribute to the crosstalk.

| $X$ | $Y$ | $\vert X\vert $ | $\vert Y\vert $ | $\ \mathbf{u}_X\ $ | $\ \mathbf{u}_Y\ $ | $\delta L_{XY}$ | $L_{XY}$ | $c$ | $r_c$ | $\rho_A$ | $\rho_u$ | $p$ | $s$ |
| --- | --- | --- | --- | --- | --- | --- | --- | --- | --- | --- | --- | --- | --- |
| ALLOGRAFT REJECTION<br>(*RPS19*, *CDKN2A*, *RPL39*) | MYC TARGETS V1<br>(*FBL*, *MYC*, *PABPC1*, *RPL18*,<br>*RPL6*, *RPLP0*, *RPS10*, *RPS2*,<br>*RPS3*, *RPS5*, *RPS6*) | 62 | 53 | 0.338 | 1.334 | 19 | 101 | 0.252 | 0.188 | 0.001 | 0.001 | 1.48E-05 | 1.219 |
| ESTROGEN RESPONSE<br>EARLY<br>(*CCND1*, *MYC*, *KRT15*) | P53 PATHWAY<br>(*CDKN2A*, *CDKN2B*, *KRT17*) | 44 | 50 | 0.593 | 0.404 | 5 | 19 | 0.120 | 0.263 | 0.001 | 0.001 | 1.48E-05 | 0.578 |
| KRAS SIGNALING DN<br>(*KRT15*, *KRT5*, *PKP1*) | P53 PATHWAY<br>(*KRT17*, *SERPINB5*) | 14 | 53 | 0.584 | 0.304 | 3 | 4 | 0.110 | 0.750 | 0.001 | 0.001 | 1.48E-05 | 0.532 |
| ANDROGEN RESPONSE<br>(*DBI*, *KRT19*, *KRT8*) | APOPTOSIS<br>(*APP*, *CLU*, *KRT18*) | 29 | 56 | 0.665 | 0.395 | 4 | 7 | 0.109 | 0.571 | 0.001 | 0.001 | 1.48E-05 | 0.528 |
| MYC TARGETS V1<br>(*MYC*, *FBL*, *RPL6*, *RPLP0*,<br>*RPS10*, *RPS2*, *RPS3*, *RPS5*,<br>*RPS6*) | P53 PATHWAY<br>(*CDKN2A*, *CDKN2B*, *RPS12*) | 53 | 52 | 1.155 | 0.292 | 10 | 27 | 0.116 | 0.370 | 0.002 | 0.001 | 2.82E-05 | 0.526 |

As a proof-of-concept, we applied our approach to study the alteration of pathway crosstalks and cell-cell communications in publicly available single-cell RNA expression data from a recent study that proposed a high-resolution map of cell diversity in normal and cancerous human breast^10,11^.

## Results

### Crosstalk quantification, statistical assessment and key players

Here we describe how we define a “crosstalk”, that is, intra- and inter-cellular interactions between two gene sets, the assessment of its statistical significance and, lastly, how we use the results of crosstalk analysis to score genes based on their contribution (**Figure 1**, see **Supplementary notes** for further details). Note that we adopt a “gene-centric” view of molecular interactions – like in gene-centric human interactomes^4,5^ – where the term “genegene interaction” refers to various types of molecular interactions (protein-protein, protein-RNA, protein-DNA) that involve the considered gene pair.

**Figure 1.**
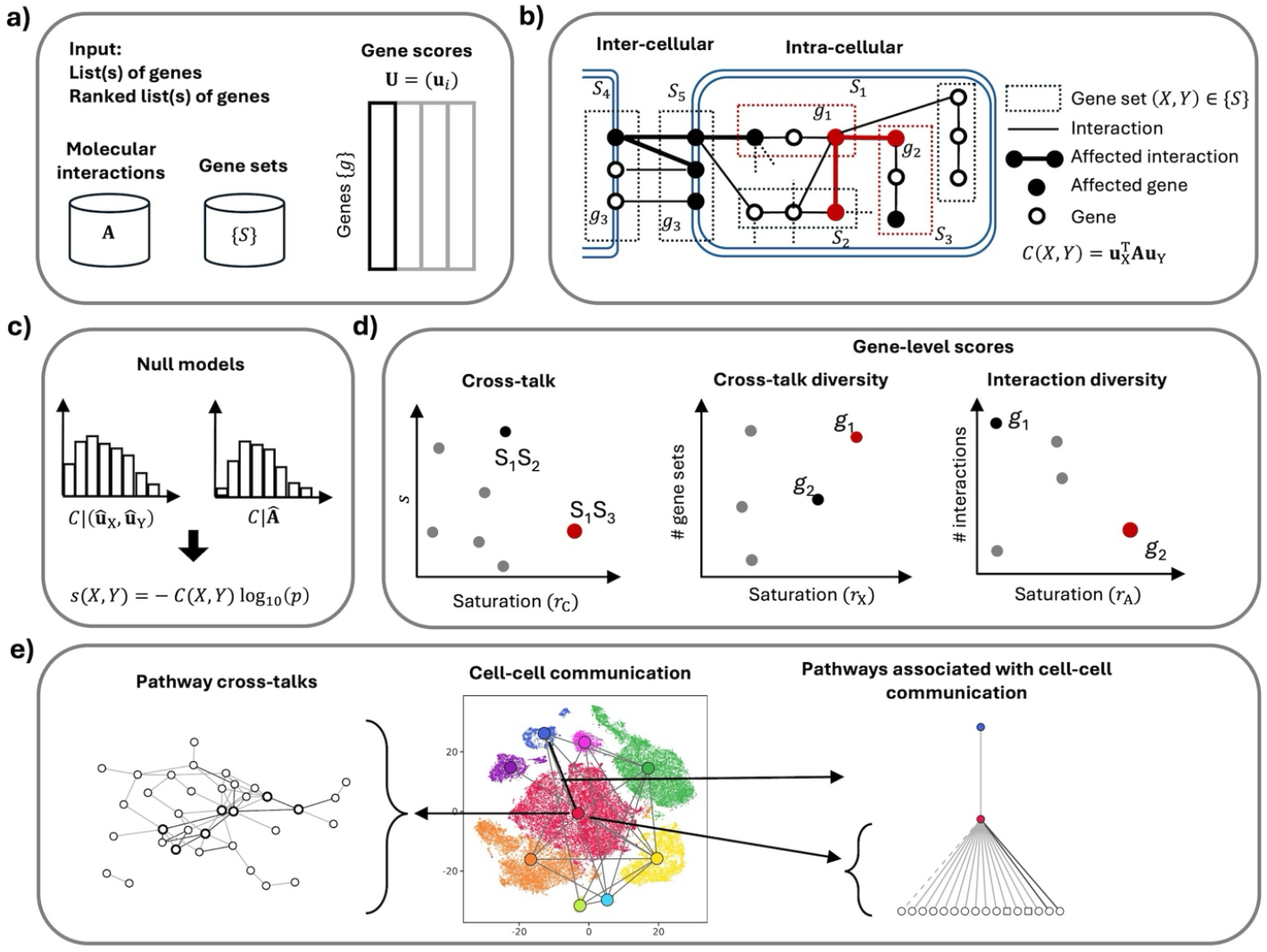
Overview of crosstalk analysis. **a)** Input data. **b)** Visualization of the molecular interactions among gene sets, at inter-cellular and intra-cellular levels, which lead to altered crosstalks. **c)** The crosstalk value is supported by two null models. **d)** Crosstalk values can be distinguished based on their saturation; the crosstalk diversity and interaction diversity are two gene-level scores that enable the identification of key crosstalk mediators; these scores can be distinguished based on their saturation. **e)** Crosstalk analysis identifies networks of intra-cellular processes, cell-cell communication and intra-cellular processes associated with cell-cell communication.

Two types of input are needed to calculate the crosstalk between any two gene sets *X* and *Y*:

- a list of gene-gene interactions, which can be derived from publicly available resources (like STRING^12^ and Omnipath^3,13^);
- one or more sets of gene-level weights (in the unit interval), which provide a summary of the gene-level alterations of interest (e.g., differential expression or mutations). Formally, we quantify the crosstalk score as the sum of weighted products between the genes of *X* that interact with those of *Y*:

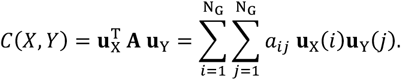

where **A** = (*a*_*ij*_) is the adjacency matrix that specifies the interactions among the N_G_ genes, while **u**_X_ and **u**_Y_ are vectors of gene weights with positive values only for the genes of *X, Y*.

The definition of the quantities involved in the calculation *C*(*X, Y*) follows three scenarios, based on whether the crosstalk is between gene sets associated with intra-cellular states, inter-cellular states or both. To quantify the alteration of crosstalk between two *intra-cellular* pathways (or another type of gene system), we exclude the genes shared between them, that is *X* ∩ *Y* = ∅, otherwise we would consider intra-pathway interactions. Further, gene weights **u**_X_ and **u**_Y_ are defined from the same input source (e.g., same set of alterations), because the focus is on an intra-cellular characteristic, and the two gene sets represent internal states of the same cell. The molecular interactions are collected from “general purpose” database of interactions, like STRING^12^.

Conversely, to quantify the *inter-cellular* crosstalk between two gene sets that are associated with two cell types, it is expected to have *X* ∩ *Y* ≠ ∅, because the two cell types, for example, can express a series of genes in common. Moreover, **u**_X_ and **u**_Y_ come from different sources, because the alterations are relative to distinct cell types. The molecular interactions are collected from databases that focus on ligand-receptor, like Omnipath^3,13^, a collection composed by multiple sources (i.e., Ramilowski^14^, CellPhoneBD^15^)

Lastly (third scenario), to quantify the intra-cellular alterations associated with inter-cellular alterations, we consider – for each cell type under analysis – the genes (set *X*) involved in any of the inter-cellular crosstalks of the cell type, and any altered pathway (set *Y*) of the cell type. Besides such peculiarity in the definition of *X* and *Y*, we have, like in the first scenario that *X* ∩ *Y* = ∅, and **u**_X_ and **u**_Y_ defined from the same input source, because they are relative to the same cell type.

A meaningful quantity that complements *C*(*X, Y*) is the crosstalk *saturation*

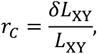

which captures, in the process under study, the number of altered interactions *δL*_XY_ between *X* and *Y*, in relation to all the possible interactions *L*_XY_ between *X* and *Y*. Indeed, similar values of *C*(*X, Y*) can be due to a higher or lower impairment of the links between the two gene sets.

To statistically benchmark the magnitude of an observed crosstalk value *c* we have to consider that it might depend on various features like gene set size, distribution of gene weights and gene degree. We focused on two null models, namely *M*_A_and *M*_u_, in both of which we preserve gene set size, degree sequence, and the association between gene weight and gene degree (within the same bin over the degree sequence), and randomize, respectively, gene-gene interactions and gene weights. Null model *M*_A_ is designed to test the dependence of *c* from the network proximity of *X* and *Y*, while *M*_u_ is meant to test the dependence of *c* from the weights of *X* and *Y* genes. This leads to four possible outcomes, determined by the possible significance of *c* in a single null, in both or in neither of them (**Supplementary Figure 1, Supplementary Table 2**). As expected by the fact that the two nulls disrupt different features, the analyses performed in our proof-of-concept (see next sections) revealed negligible correlations between the values obtained with the two nulls (**Supplementary Figure 2**). It is therefore meaningful to combine the two probabilities *ρ*_A_ and *ρ*_u_ of observing – respectively – a value equal or greater than *c* in *M*_A_ and *M*_u_, into the probability of observing a product 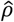 as small as the one observed

**Figure 2.**
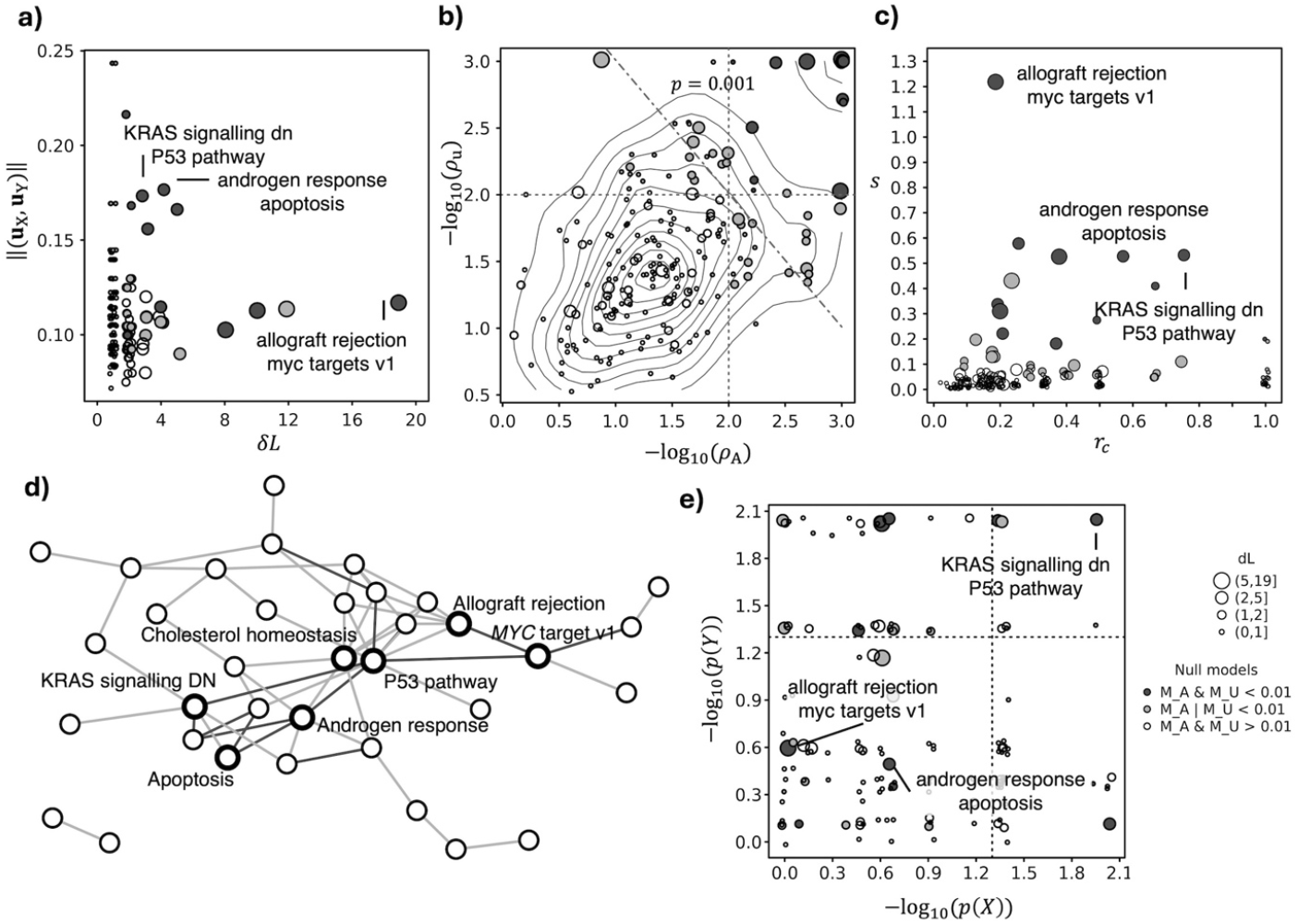
Intra-cellular crosstalks controlled by expression changes specific of TNBC cells. **a)** number of altered links (*δL*) and average gene weight (‖(**u**_X_, **u**_Y_)‖) of the crosstalk forming genes. **b)** The probabilities *ρ*_A_ and *ρ*_u_ estimated by the two null models for each crosstalk value; the vertical and horizontal lines denote *α* = 0.01, while the diagonal line denotes *p* = 0.001. **c)** Crosstalk score *s* and its saturation *r*_*c*_. **d)** Network of processes that establish crosstalks supported (*α* = 0.01) by at least a null model. **e)** Over representation analysis p-values *p*(*X*) and *p*(*Y*) for each of the processes (*X, Y*) that establish a crosstalk; the vertical and horizontal lines denote *α* = 0.05.

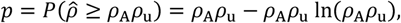

which is equal to the probability given by means of the so-called Fisher’s combined probability test ^16,17^.

Lastly, we define a summary score for ranking crosstalks, combining effect size *C*(*X, Y*) and its estimated probability *p*:

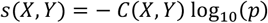

The list of significant crosstalks provides the opportunity to score genes based on their contribution. We consider two gene-level quantities: *crosstalk diversity* and *interaction diversity*. The first counts how many gene sets that are part of altered crosstalks contain interactors of a gene *g*_*i*_; therefore, the saturation *r*_X_(*i*) of the crosstalk diversity of *g*_*i*_ reaches 1 when all the gene sets that contain interactors of *g*_*i*_ are part of altered crosstalks. The second counts the interactors of *g*_*i*_ that belong to gene sets that are part of crosstalks; analogously to *r*_X_(*i*), the saturation *r*_A_(*i*) of the interactor diversity of *g*_*i*_ reaches 1 when all the interactors of *g*_*i*_ belong to gene sets that are part of altered crosstalks.

### Alteration of crosstalks in triple negative breast cancer

As a proof-of-concept, we analysed the crosstalks in triple negative breast cancer (TNBC), using single-cell RNA expression data from a recent study that proposed a high-resolution map of cell diversity in normal and cancerous human breast^10,11^. Our objective is to show what kind of information can be extracted from the analysis of crosstalks using data generated by means of one the state-of-the-art technologies in transcriptomic analysis. In particular, we focused on cancer cells and analysed the interactions among intra-cellular processes whose alteration could be implied in the dysregulation of the reciprocal control among molecular mechanisms that could contribute to tumour progression. Then, we considered the communication between cancer cells and Cancer Associated Fibroblasts (CAF). Indeed, CAFs represent a peculiar hub of cell-cell communication within the tumour niche by promoting tumoral growth and malignancy by releasing factors targeting cancer cells, repressing immune response by their interactions with immune cells and inducing angiogenesis interacting with endothelial cells^18–20^. Lastly, we shed light to cancer cell processes that could be associated with the communication between CAFs and cancer cells.

### Alteration of intra-cellular crosstalks in cancer cells

We screened the impact of 304 (**Supplementary Tables 3-4**) cancer epithelial cell markers (*p* < 0.05, log.(FC) > 0.5, Cancer epithelial vs all) on the crosstalk between intra-cellular processes (MSigDB Hallmarks database^21^). We found that most of the crosstalks is altered in up to 2 interactions and that the maximum number of altered interactions is 19, among a total of 14 genes belonging to allograft rejection and *MYC* targets (v1) (**Table 1, Figure 2a, Supplementary Table 5**). We observed a marked variability of gene weights, degree of statistical significance, and saturation, independently from the number of affected links, which makes such pieces of information useful to differentiate crosstalks (**Figure 2a-c**). In particular, we observed a total of 59 crosstalks whose score can hardly be obtained (*α* = 0.01) when shuffling interactions or gene weights, and 14 crosstalks that are supported by both nulls (**Figure 2b**). Among these, we obtained several crosstalks that involves the p53 pathway, cholesterol homeostasis and androgen response, which emerge as hubs in the network of altered crosstalks **(Figure 2d)**. In 24 crosstalks, the saturation indicates the alteration of more than half of the links (**Figure 2c**), like between p53 pathway and KRAS signalling (“KRAS_SIGNALING_DN”), which involves the alterations of 3 out of 4 interactions between a total of 5 genes, and the score is supported by both nulls (**Figure 2c, Supplementary Table 5)**. To compare the outcomes of crosstalk and pathway enrichment analyses, we assessed to which extent the processes exhibiting significant crosstalks are also marked by significant enrichment in DEGs (**Figure 2d, Supplementary Table 6**). As expected, the two types of analyses provide a complementary view, where several processes involved with significant crosstalk do not display enrichment and *vice versa*. Only in a few cases (7 pairs) both the processes are enriched (*p*-value < 0.05) in DEGs, while the majority of altered crosstalks takes place between pairs of processes that are not enriched in DEGs.

Significantly altered crosstalks are mediated by a total of 57 genes (**Supplementary Table 7**). Crosstalk diversity and interaction diversity suggest a gene prioritization that is independent from their initial alteration score. In other words, genes that were ranked low by differential expression analysis can emerge as key players as mediators of crosstalks. This is the case of the two cyclin-dependent kinases *CDKN2A* and *CDKN2B*, which stand out for their crosstalk diversity, as they mediate 11 and 10 altered crosstalks, respectively (**Figure 3a**). Among the genes with the highest interactor diversity we obtained a series of genes that code for ribosomal-associated proteins (**Figure 3b**). We observed a wide range of saturations and an overall correlation between the saturation of crosstalk diversity and that of interaction diversity **(Supplementary Table 7)**. Among the genes with the highest values of both saturations we found the two tumour proteins D52 (*TPD52*) and D53 (also known as *TPD52L1*) (**Figure 3**), which are involved in cancer cells proliferation and more aggressive phenotype^22,23^.

**Figure 3.**
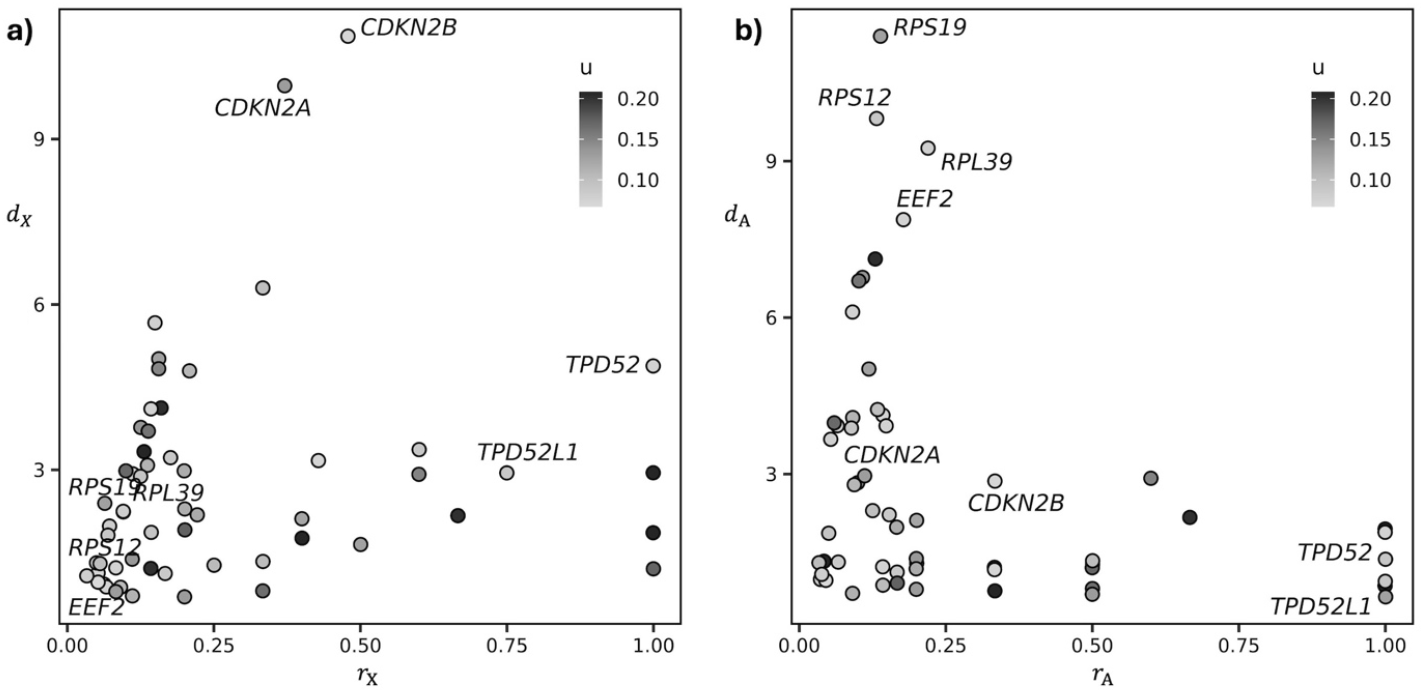
Crosstalk diversity and interaction diversity of the DEGs that are involved in intra-cellular crosstalks in TNBC cells. **a)** Crosstalk diversity *d*_X_ and its saturation *r*_X_. **b)** Interactor diversity *d*_A_ and its saturation *r*_A_.

### Inter-cellular crosstalks (cell-cell communication)

We analysed the inter-cellular interactions (Omnipath^3^) among the 36 pairs of sets defined by the differentially expressed genes (FDR < 0.05, log_2_(FC) > 0.5) of 9 cell types in relation to the others, in 8 TNBC tumors^10^ (**Supplementary Tables 3-4**). Compared to the intracellular crosstalks among processes, here we dealt with larger gene sets and more links among them. As expected, this scenario led to a higher number of altered interactions, with a median of 38 and a maximum of 162 between CAFs and endothelial cells (**Figure 4a, Supplementary Table 8**). Further, the crosstalks are statistically supported (*α* = 0.01) mostly by their gene weights (21 pairs), rather than interactions, which support 3 crosstalks that are supported by both nulls, namely between B cells and tumour-associated macrophage (TAMs), between dendritic cells (DCs) and TAMs, and between B cells and DCs. Saturation reaches up to one quarter of the possible interactions between CAFs and endothelial cells (**Figure 4b**). The emerging cell-cell communication network (**Figure 4c**) highlights a relevant role of those microenvironment cells, which establish several significant interactions. The communication between cancer cells and CAFs is supported (*α* = 0.01) by randomization of gene weights, and involves 22 interactions between a total of 33 DEGs (**Figure 4c, Supplementary Table 9**). Among the key players of this communication, we found *MDK* and *MFGE8* (expressed in cancer cells), which mediate 4 and 3 interactions, respectively, with genes expressed in CAFs, including integrins *ITGB1* and *ITGB5* (**Supplementary Table 9**).

**Figure 4.**
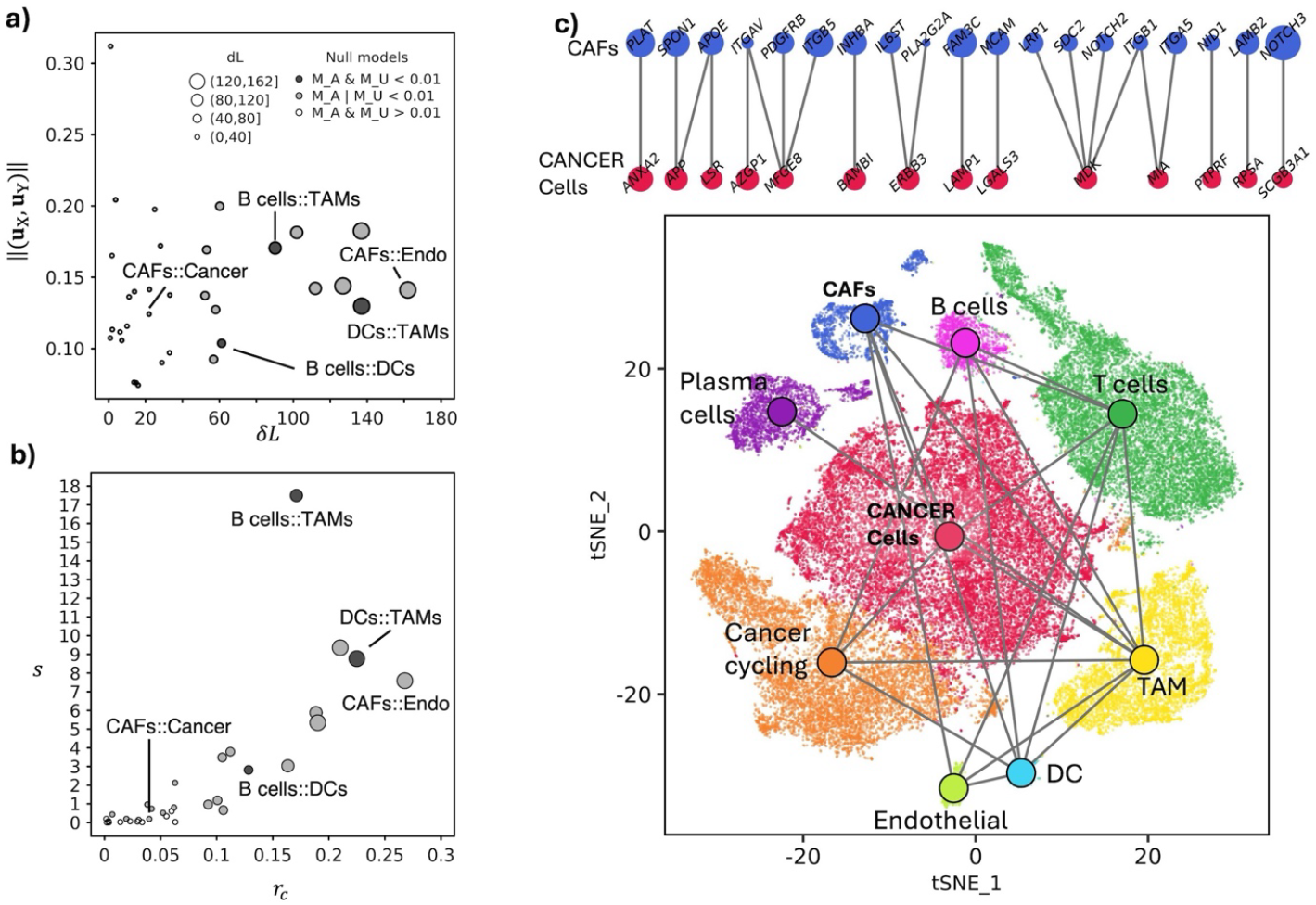
Inter-cellular crosstalks controlled by expression changes in the 9 cell types of TNBC samples. **a)** Number of altered links (*δL*) and average gene weight (‖(**u**_X_, **u**_Y_)‖) of the crosstalk forming genes. **b)** Crosstalk score *s* and its saturation *r*_*c*_. **c)** Above: DEGs that mediate the communication between CAFs and cancer cells; below: the position of cells in the space of the first two tSNE dimensions (bottom), coloured by cell type whose communications supported (*α* = 0.01) by at least a null model are indicated through a link between the two centroids.

The cell-cell communication network (*α* = 0.01) involves 379 genes (**Figure 5, Supplementary Table 10**). Among the genes that stand out for their ubiquity we observed *CXCR4*, with a crosstalk diversity of 8 (out of 9 cell-types present), and *ICAM1, TGFB1, ITGB2, PTPN6* and some Major Histocompatibility Complex genes (*HLA-C, HLA-DRA, HLA-DRB1*), which show a crosstalk diversity of 7. Among the 29 DEGs in cancer epithelial cells (out of 379), *MFGE8, LAMP1, RPSA* and *AZGP1* are specific (*d*_X_ = 1, *r*_X_ = 1) of the communication with CAFs **(Figure 5)**. Conversely, we did not observe DEGs in CAFs that are specific to the signalling with cancer cells. However, there is one gene, namely *PLAT*, which is only involved in the communication with cancer cells (*d*_X_ = 1).

**Figure 5.**
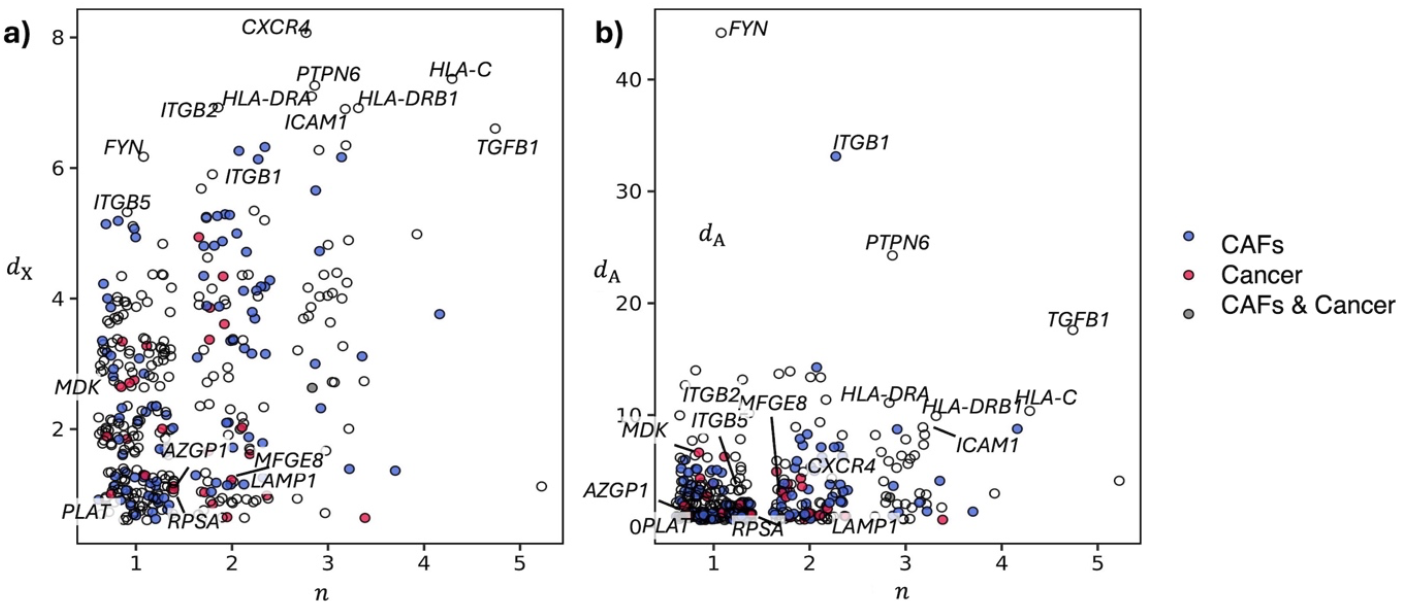
Crosstalk diversity and interaction diversity of the DEGs that are involved in inter-cellular crosstalks between CAFs and cancer cells. **a-b)** Crosstalk diversity *d*_X_ **(a)** and Interactor diversity *d*_A_ **(b)** in relation to the number of cell types (*n*) in which the gene is differentially expressed.

### Integrated crosstalks: cancer cell pathways that can be associated with the communication between cancer cell and CAFs

We analysed the crosstalks between the gene set of the 14 cancer cell DEGs that mediate the communication with CAFs, and all the processes (MSigDB Hallmarks) that contain cancer cell DEGs (**Figure 6, Supplementary Table 11**). We found 15 interactions supported (*α* = 0.01) by at least a null model and two, supported by both nulls. The first involves interactions among *RPSA*, which mediate the interaction with CAFs, and other ribosomal proteins (*RPL18, RPL6, RPLP0, RPS10, RPS2, RPS3, RPS5, RPS6*) that are regulated by *MYC*. The second take place between, on the one hand, *APP* and *PTPRF* (mediators of the communication with CAFs), and, on the other hand, *CLU* and *CTNNB1* (cholesterol homeostasis). Almost all the interactions found (13 out of 15) involve processes that establish significant (*α* = 0.01) intra-cellular crosstalks (**Supplementary Table 5**). The two processes that did not emerge in the screening of intra-cellular crosstalks of cancer cells (namely complement and coagulation) are both mediated by the interaction between *APP* and *CLU*.

**Figure 6.**
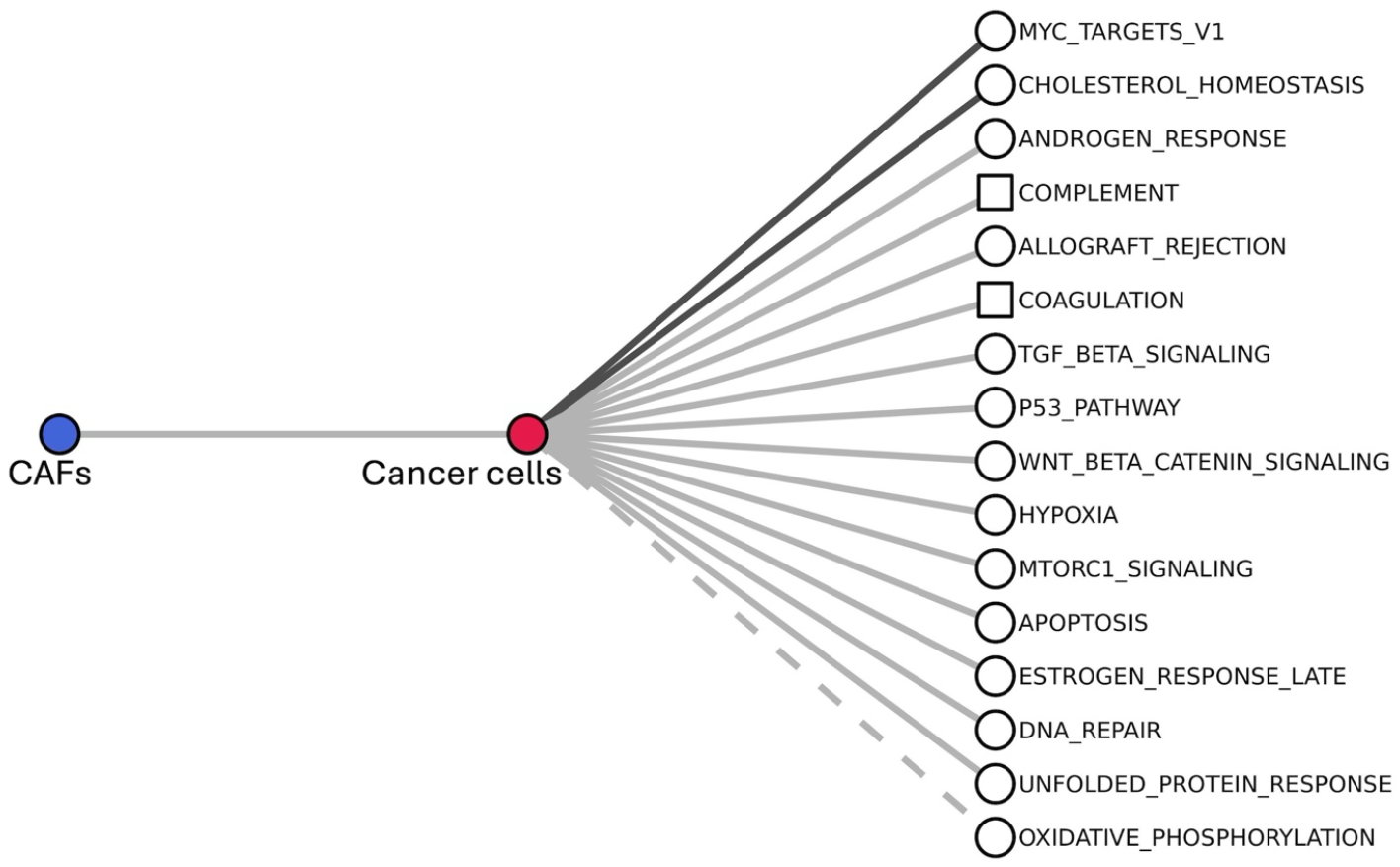
Intra-cellular processes of cancer cells associated with their signalling with CAFs. T The processes are ranked from top to bottom by decreasing value of *s*; squares indicate processes that were not found in the analysis of intra-cellular crosstalks; link colour indicates statistical evidence (as in Figures 2, 4), with the exception that, here, the dashed line replaces the white colour in indicating that both null models are above *α* = 0.01.

## DISCUSSION

We presented a network-based approach to assess the alterations of crosstalks between gene sets, based on gene-centric molecular interactions and one or more lists of gene scores that result from omics data analysis. The approach can be applied to inter-cellular as well as intra-cellular crosstalks, and to the analysis of intra-cellular crosstalks that can be associated with inter-cellular crosstalks. As a proof-of-concept, we applied the approach to analyse the crosstalks affected by the gene expression alterations detected at single-cell resolution in triple negative breast cancer samples.

The score of a crosstalk is proportional to the interactions between two gene sets and the weights of the interacting genes. The score is supported by two complementary null models that conserve gene set size, degree sequence, and the association between gene weight and gene degree. These nulls provide a means to assess whether the statistical significance of the score comes from gene weights, interactions or both. In the proof-of-concept, we showed that all three scenarios emerge when using real data.

We reported altered crosstalks at various degree of saturation, especially in the analysis of intra-cellular crosstalks. This quantity enabled the identification of pairs of processes where most of the interactions involved gene expression changes or, on the opposite, pairs of processes where only a specific part of their interaction is affected. For example, our analysis identified that the pathway of KRAS is involved in crosstalk dysregulation associated with TNBC, supporting evidence that indicates this pathway as crucial in phenotypical and metabolic features of cancer cells^24^.

We showed that the analysis of intra-cellular crosstalks complements the typical enrichment analysis. Indeed, we reported a series of gene sets that, despite not showing significant enrichments in DEGs, were part of significantly altered crosstalks. This is the case of one of the top ranked crosstalks (supported by both nulls), which suggests the impairment of regulative mechanisms between androgen response and apoptosis. Notably, the relation between androgen receptor and apoptosis has been implicated in breast cancer metastasis^25,26^. Another example is cholesterol homeostasis, which emerged as a hub of the intra-cellular network and is reported to promote cancer cell proliferation in TNBCs^27^.

The reconstruction of inter-cellular communications based on cell type-associated gene sets provides a means to overcome the heterogeneity at gene expression level and sheds light on the general picture of the active (or altered) communications among the cell types. The analysis of TNBC cell types confirmed the well-known core network of communications between cancer cells and microenvironment. Cancer cells show significant communication with CAFs, supporting the pro-tumoral role of CAFs by activating the signalling associated with proliferation and tumour progression^18–20,28^.

The joint analysis of intra-cellular and inter-cellular cross talks paves the way towards the reconstruction of maps that integrate the communications between different cell types with the pathway crosstalks activated within each one. In the proof-of-concept we analysed the processes that are activated in cancer cells and can be associated with their communication with CAFs. Notably, a mediator of such crosstalk is the extracellular chaperone *CLU*, which was reported as a key player in cancer^29^ and an interesting actionable target in TNBC^30,31^.

With the aim of providing an additional way of identifying the genes associated with a phenotype, we introduced the crosstalk diversity and interaction diversity. These quantities shed light on the genes that act as mediators of the signalling between processes or cell types. We showed, as a proof-of concept, that a series of genes with extreme crosstalk diversity and interaction diversity is indeed known to be associated with the process under study. Among them, *EEF2* was demonstrated to be upregulated in several cancers and associated with worse prognosis, thus suggesting its potentiality as novel therapeutic target^32,33^. The high interaction diversity of ribosomal-associated proteins sustains the importance of the dysregulation of translation process in tumorigenesis mechanisms and the clinical potential represented by targeting this process in tumour cells^34^. Interestingly, some of the genes prioritized by crosstalk diversity and interactor diversity have marginal expression changes and, therefore, stand out due to their pattern of interactions with other altered genes. This is the case of *CDKN2A* and *CDKN2B*, which exert a role in the regulation of cell cycle and proliferation and their association with breast cancer is largely studied^35,36^. The analysis of genes that mediate the inter-cellular communications revealed a series of genes shared by multiple communications. These genes are involved in tumour promoting functions supporting tumour growth, chronic inflammation and angiogenesis, by secretion of growth factors and other soluble molecules, vesicles, and mechanic interactions among cells and extracellular matrix^20,37–39^. Concerning the genes that mediate the signalling between CAFs and cancer cells, *MDK* and *MFGE8* (expressed in cancer cells) are known to be associated with the acquisition of various tumour hallmarks^40,41^. Studies suggest the involvement of *AZGP1* in the differentiation of progenitor cells into CAF to support tumorigenesis^42^, while *RPSA* and *LAMP1* are implicated in poor prognosis in breast cancer^43,44^. *PLAT* was reported to regulate the ability of breast cancer CAFs to invade stroma^45^, and as an angiogenetic factor of CAF associated with negative prognosis in colon cancer^46^.

Crosstalk diversity and interaction diversity can be relevant for the choice of actionable targets. Genes that affect several crosstalks interacting with multiple cellular functions are interesting targets for therapy, but – at the same time – could be associated with a wide spectrum of negative side effects. The saturations of crosstalk diversity and interaction diversity provide a means to collect more selective targets for therapy, as it prioritizes genes that mediate crosstalks with less but more disease-specific cellular functions.

The results presented in this study have to be seen in light of some limitations. The gene-gene interactions available in the literature are aspecific, and as such, they are a model of the interactions that *potentially* take place in the biological system under analysis. Moreover, the collections of molecular interactions are known to be affected by the various biases^4,5^. We have used state-of-the art collections and filtered the interactions to ensure an appropriate trade-off between coverage of genes and presence of biases, following the recommendations of previous studies^4,5^. As a proof-of-concept, we studied the intra-cellular crosstalks using gene set definitions from MSigDB hallmarks. There are multiple ways to define intra-cellular processes, e.g. using databases like KEGG and Reactome. Therefore, other analyses of intra-cellular crosstalks in cancer cells of TNBC are possible and could highlight additional mechanisms. To perform the proof-of-principle, we considered scRNA sequencing data from a recent study in breast, which allowed us to analyse intra-cellular as well as inter-cellular crosstalks, and the two of them combined. However, the number of tested genes was limited by the sensitivity and depth of the type of technology used in such study. In turns, the results emerged in the proof-of-principle should be interpreted considering this limited observability of the underlying real processes.

In conclusion, the approach presented in this work and the results gained in the proof-of-principle, even in the light of their limitations, support the usefulness of crosstalk analysis as an additional instrument to the “toolkit” of biomedical research for translating complex biological data into actionable insights.

## Methods

### Definition of gene weights from single cell RNA-sequencing data

The Seurat data object “SeuratObject_TNBC.rds” containing single-cell RNA expression data of 8 triple negative breast tumors^10,11^ was downloaded from figshare^47^. The associations between the 9 cell clusters and cell types (not available in the Seurat object) were obtained on the basis of the cell association provided by the authors in the figures of the paper, together with “SeuratObject_TNBCSub.rds” object **(Supplementary Figure 3)**. Differentially expressed genes were obtained by means of MAST algorithm^48^, testing each cell type against all the other cells (Seurat^49^ function “FindAllMarkers()”, default parameters). Differential expression statistics were used to define a gene weight vector **u**_*j*_ (of size equal to the total number of genes in the considered analysis) for each cell type *j* combining log fold change *x* and adjusted *p*-value (Benjamini-Hochberg method^50^); to reduce noise, scores associated with marginal significance were set to zero, that is: *y*_*ij*_ = ™log_2_(*x*) log_10_(*p*), when *p* < 0.05 ∧ log_2_(*x*) ≥ 0.5, while *y*_*ij*_ = 0 otherwise, where *i* is the index for genes. Each vector was normalized to have a maximum value of 1: *u*_*ij*_ = *y*_*ij*_/max_*i*_(*y*_*ij*_).

### Molecular interactions and gene sets

Molecular interactions used for pathway crosstalk analysis were downloaded from STRING^12^ (v12, https://string-db.org/cgi/download). The combined score was updated excluding “text mining” using a modified version of the script “combine_subscores.v2.py” (https://stringdb-downloads.org/download). Ensembl identifiers were mapped to Entrez Gene identifiers using the mapping available in STRING (https://string-db.org/cgi/download) and Entrez Gene (ftp://ftp.ncbi.nih.gov/gene/DATA, September, 19, 2023). The highest score was considered for each gene pair. Only high-confidence (combined score ≥ 700) interactions and the top 3 (per gene) interactions with medium confidence (STRING score ≥ 400) were considered, obtaining a total of 174’962 interactions involving 17’288 genes. Molecular interactions available in Omnipath^13^ were obtained through the R package OmnipathR^51^ (September, 2024), for a total of 4’312 interactions involving 1’782 genes. The MSigDB Hallmarks gene sets^21^ were collected through the R package “msigdbr” v7.4.1^52^.

In each analysis, the initial gene set list was created to ensure that: each gene had at least an interaction; only gene sets with at least 3 elements and a non-null gene weight were considered; to reduce the number of possible gene set pairs, only those such that *C*(*X, Y*) > 0 were considered.

### Randomizations and computational aspects

A total of 1000 randomizations of gene labels was used to create the null models. Gene degree was preserved splitting the degree sequence in equally sized bins, 9 for intra-cellular crosstalks, 4 to study inter-cellular communications, and 7 to study cancer cell intracellular crosstalks associated with their communication with CAFs). The number of bins was optimized to use the highest number, between 2 and 15, that leads to non-empty intervals. The average computational cost for the analysis of intracellular crosstalks with 203 gene set pairs was approximately 4 minutes over 8 cores with 64GB of RAM per core.

## Code availability

The computational method used in this study (Ulisse v2.0) is available in Zenodo with the identifier 10.5281/zenodo.15166722. Source code and documentation are freely available in github at the URLs https://github.com/emosca-cnr/Ulisse and https://emosca-cnr.github.io/Ulisse.

## Data availability

The single-cell RNA sequencing data that support the findings of this study are available in “figshare” with the identifier 10.6084/m9.figshare.17058077.v1^47^.

## Supplementary Information

Supplementary notes, figures and tables are available in the files “Supplementary_Information.pdf” and “SupplementaryTables_3-11.xlsx”.

## Funding

This work was supported by Fondazione Regionale per la Ricerca Biomedica (Regione Lombardia) (ERAPERMED2018-233 FindingMS GA 779282), European Commission H2020 GEMMA project (ID 825033), and European Union NextGenerationEU, Mission 4 Component 2, project “Strengthening BBMRI.it”, CUP B53C22001820006. The funders had no role in study design, data collection and analysis, decision to publish, or preparation of the manuscript.

## Notes

### Competing Interest Statement

The authors have declared no competing interest.

### Summary of Updates

- crosstalk quantification formalism; - null models; - proof-of-concept: datasets and analysis; - main text, figures, tables and supplementary information.

