## Supplementary for "Analysis of intracellular and intercellular crosstalk from omics data"

*Corresponding author

### Supplementary Notes

#### Definition of crosstalk and its saturation

Here, we describe how we quantify a "crosstalk" between two gene sets. We consider the general scenario where input data derive from single-cell RNA-sequencing experiments, because this enables the calculation of both intra-cellular and extra-cellular crosstalks. In this case we have input vectors $(\mathbf{u}_1, \dots, \mathbf{u}_\mathrm{m})$, each of which contains gene-level scores that characterize the "profile" of a cell type (or cluster), like, differential expression scores between a cell type and the others. Let us consider:

- the universe $\Sigma = \{X_i\}_{i=1}^{\mathrm{N_S}}$ of gene sets $X$ that we want to analyze; the calculation of *intra-cellular crosstalk* require $X \cap Y = \emptyset \ \forall (X, Y) \in \Sigma^2$;
- the totality of genes, which forms the gene set $V = \bigcup_{i=1}^{\mathrm{N_S}} X_i = \{g_i\}_{i=1}^{|\mathrm{N_G}|}$ , indexed by the set $I = \{1, \dots, \mathrm{N_G}\}$;
- the undirected graph $G = (V, E)$ among the genes, and the corresponding adjacency matrix $\mathbf{A} \in \{0,1\}^{\mathrm{N_G} \times \mathrm{N_G}}$;
- the $\mathrm{N_G} \times \mathrm{m}$ weight matrix $\mathbf{U} = (\mathbf{u}_1, \dots, \mathbf{u}_\mathrm{m})$, $\mathbf{U} \in [0, 1]^{N_\mathrm{G} \times \mathrm{m}}$;
- the weight function $w$ that defines the gene weight vector associated with the gene set $X$ as: $\mathbf{u}_\mathrm{X} = w(X, j; \mathbf{U}) = \mathrm{diag}(\mathbf{1}_\mathrm{X})\ \mathbf{U}_{*j}$, where $\mathbf{1}_\mathrm{X}$ denotes the indicator vector of $X$, $\mathbf{1}_\mathrm{X}(i) = \begin{cases} 1, & i \in K_\mathrm{X} \\ 0, & i \notin K_\mathrm{X} \end{cases}$, and $K_\mathrm{X} \subseteq I$ is the set of indices of $g_i \in X$.

The crosstalk of any gene set pair $(X, Y) \in \Sigma^2$ is:

$$C(X, Y) = \mathbf{u}_\mathrm{X}^\mathrm{T}\ \mathbf{A}\ \mathbf{u}_\mathrm{Y} = \sum_{i=1}^{\mathrm{N_G}} \sum_{j=1}^{\mathrm{N_G}} a_{ij}\ \mathbf{u}_\mathrm{X}(i) \mathbf{u}_\mathrm{Y}(j).$$

Typically – as explained in the main text – the calculation of *intra-cellular crosstalk* is done using the same column of $\mathbf{U}$ for $X$ and $Y$: $\mathbf{u}_\mathrm{X} = w(X, j)$, $\mathbf{u}_\mathrm{Y} = w(Y, j)$.

Let us consider all pairs of sets $(X, Y)$ that establish a crosstalk $C(X, Y) > 0$. Let us define the number of links between $X$ and $Y$ as $L_\mathrm{XY} = \mathbf{1}_\mathrm{X}^\mathrm{T}\ \mathbf{A}\ \mathbf{1}_\mathrm{Y}$, and the number of links between $X$ and $Y$ affected by a molecular change as $\delta L_\mathrm{XY} = \mathbf{b}_\mathrm{X}^\mathrm{T}\ \mathbf{A}\ \mathbf{b}_\mathrm{Y}$, where $\mathbf{b}_\mathrm{X}(i) = \mathrm{sgn}(\mathbf{u}_\mathrm{X}(i))$ and $\mathbf{b}_\mathrm{Y}(i) = \mathrm{sgn}(\mathbf{u}_\mathrm{Y}(i))$. The saturation of the crosstalk is defined as

$$r_c = \frac{\delta L_\mathrm{XY}}{L_\mathrm{XY}}.$$

#### Statistical assessment of crosstalks

As stated in the main text, to statistically benchmark the magnitude of an observed crosstalk value $c$ we have to consider that it might depend on various features (e.g., gene set size, distribution of gene weights and gene degree). We focused on two null models, namely $M_\mathrm{A}$and $M_\mathrm{u}$, in both of which we *preserve* gene set size, degree sequence, and the association between gene weight and gene degree (within the same bin over the degree sequence), and randomize, respectively, gene-gene interactions and gene weights. Null model $M_{\mathrm{A}}$ is designed to test the dependence of $c$ from the network proximity of $X$ and $Y$, and allows us to estimate the probability of observing a value equal or greater than $c$, using a set of permutations $\widehat{\mathbf{A}} = \mathbf{PAP}$ where $\mathbf{P}$ is a (gene-degree preserving) permutation matrix:

$$\rho_{\mathrm{A}} = \Pr\big(C\big(X, Y; \widehat{\mathbf{A}}, \mathbf{u}_{\mathrm{X}}, \mathbf{u}_{\mathrm{Y}}\big) \geq c\big).$$

Null model $M_{\mathrm{u}}$ is meant to test the dependence of $c$ from the weights of $X$ and $Y$ genes and allows us to estimate the probability of observing a value equal or greater than $c$ using permutations $\widehat{\mathbf{U}} = \mathbf{PU}$

$$\rho_{\mathrm{u}} = \Pr\big((X, Y; \mathbf{A}, \widehat{\mathbf{u}}_{\mathrm{X}}, \widehat{\mathbf{u}}_{\mathrm{Y}}) \geq c\big),$$

where $\widehat{\mathbf{u}}_{\mathrm{X}}$ and $\widehat{\mathbf{u}}_{\mathrm{Y}}$ are derived from $\widehat{\mathbf{U}}$ by means of the weight function $w$.

Then, we combine the two probabilities $\rho_{\mathrm{A}}$ and $\rho_{\mathrm{u}}$ into the probability of observing a product $\hat{\rho}$ as small as the one observed, which is[1,2]:

$$p = \Pr(\hat{\rho} \geq \rho_{\mathrm{A}}\rho_{\mathrm{u}}) = \rho_{\mathrm{A}}\rho_{\mathrm{u}} + \int_{\rho_{\mathrm{A}}\rho_{\mathrm{u}}}^{1} \frac{\rho_{\mathrm{A}}\rho_{\mathrm{u}}}{\rho_{\mathrm{A}}} \mathrm{d}\rho_{\mathrm{A}} = \rho_{\mathrm{A}}\rho_{\mathrm{u}} - \rho_{\mathrm{A}}\rho_{\mathrm{u}} \ln(\rho_{\mathrm{A}}\rho_{\mathrm{u}}).$$

Note that $p$ is equal[1] to the probability estimated by the so-called Fisher's combined probability test[2]; we refer the interested readers to Wallis (1942)[1] for further details.

Lastly, we define a summary score for ranking, combining effect size $C(X, Y)$ and its estimated probability $p$:

$$s(X, Y) = -\,C(X, Y) \log_{10}(p).$$

#### Definition of crosstalk diversity and interaction diversity

Let us consider all pairs of sets $(X, Y)$ that establish a crosstalk $C(X, Y) > 0$, a subset of which is considered significant $s(X, Y) > \alpha$, given a chosen threshold $\alpha$. The list of significant, or altered, crosstalks is then used to score the genes $g_i$ that mediate such interactions. To this aim, we introduce the gene-level quantities *crosstalk diversity* and *interaction diversity*. Define *the crosstalk diversity* of $g_i$ as the number $\delta d_{\mathrm{X}}(i)$ of gene sets $Y$ such that:

- $Y$ is involved in an *altered* crosstalk with a set $X$ that contains $g_i$;
- $Y$ contains a gene $g_j$ that mediates the crosstalk interacting with $g_i$.

Formally:

$$\delta d_{\mathrm{X}}(i) = \big|\{Y \mid g_i \in X, g_j \in Y, a_{ij}\, \mathbf{u}_{\mathrm{X}}(i)\, \mathbf{u}_{\mathrm{Y}}(j) > 0, s(X, Y) > \alpha\}\big|.$$

We normalize this quantity considering the gene sets $Y$, such that:

- $Y$ is involved in a crosstalk with a set $X$ that contains $g_i$;
- $Y$ contains a gene $g_j$ that interacts with $g_i$.

That is:

$$d_{\mathrm{X}}(i) = \big|\{Y \mid g_i \in X, g_j \in Y, a_{ij} = 1, C(X, Y) > 0\}\big|.$$

The ratio of the two numbers above is the *saturation* of the crosstalk diversity of $g_i$:

$$r_{\mathrm{X}}(i) = \frac{\delta d_{\mathrm{X}}(i)}{d_{\mathrm{X}}(i)}.$$

Define *interaction diversity* of $g_i$ as the number $\delta d_{\mathrm{A}}(i)$ of genes $g_j$, such that:

- $g_j$ belongs to $Y$, a set that establishes an *altered* crosstalk with $X$, which contains $g_i$;

- $g_j$ interacts with $g_i$.

Formally:

$$\delta d_{\mathrm{A}}(i) = \left|\left\{g_j \mid g_j \in Y, g_i \in X, a_{ij}\, \mathbf{u}_{\mathrm{X}}(i)\, \mathbf{u}_{\mathrm{Y}}(j) > 0, s(X,Y) > \alpha\right\}\right|.$$

We normalize this quantity considering the genes $g_j$, such that:

- $g_j$ belongs to $Y$, a set that establishes a crosstalk with $X$, which contains $g_i$;
- $g_j$ interacts with $g_i$.

Formally:

$$d_{\mathrm{A}}(i) = \left|\left\{g_j \mid g_j \in Y, g_i \in X, a_{ij} = 1, C(X,Y) > 0\right\}\right|.$$

The *saturation* of the interaction diversity of $g_i$ is:

$$r_{\mathrm{A}}(i) = \frac{\delta d_{\mathrm{A}}(i)}{d_{\mathrm{A}}(i)}.$$

### Supplementary Figures

**Supplementary Figure 1**. Gene weights, gene-gene interactions and crosstalk values of four gene sets pairs that exemplify the four combinations of evidence about the statistical significance of a crosstalk based on the two null models $M_{\mathrm{A}}$ and $M_{\mathrm{u}}$.

**a-d).** Average weight $|\mathbf{u}|$ (horizontal axis) of all the genes that are involved in the crosstalk between a pair of gene sets $X$ and $Y$, and number of interactions $|\mathbf{A}|$ (vertical axis) between the genes of $X$ and those of $Y$, in 1000 degree-preserving permutations of gene labels applied to gene weights (null model $M_{\mathrm{u}}$) or gene-gene interactions (null model $M_{\mathrm{A}}$). **e-h).** Crosstalk values in null models $M_{\mathrm{u}}$(horizontal axis) and $M_{\mathrm{A}}$ (vertical axis) for the same gene sets pairs and permutations shown in panels a-d. **a-h)** The analysis was performed as described in the main text section "Alteration of intra-cellular crosstalks in cancer active fibroblasts". **a,e)** (OXIDATIVE_PHOSPHORYLATION, P53_PATHWAY): not significant in any null. **b,f)** (E2F_TARGETS, G2M_CHECKPOINT): significant in $M_{\mathrm{A}}$; **c,f)** (MYC_TARGETS_V1, OXIDATIVE_PHOSPHORYLATION): significant in $M_{\mathrm{u}}$; **d,h)** (ALLOGRAFT_REJECTION, MYC_TARGETS_V1): significant in both nulls.

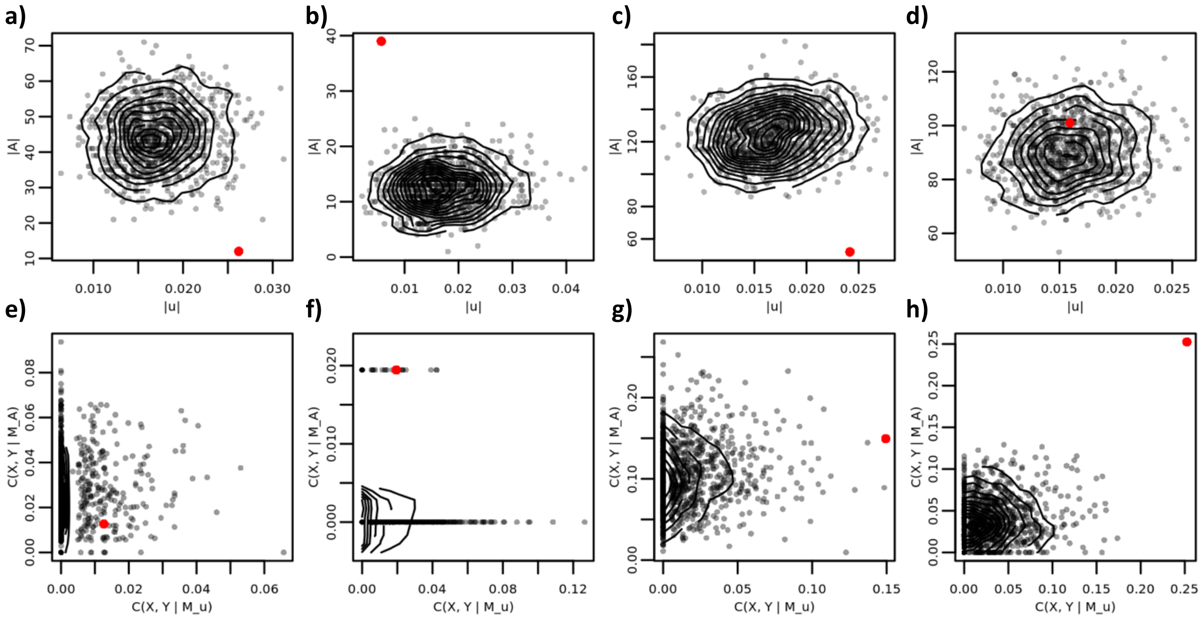

**Supplementary Figure 2**. Frequency distribution of the Spearman correlation values between the two null models $M_{\mathrm{A}}$ and $M_{\mathrm{u}}$.

The correlations shown are between crosstalk values (1000 permutations) in the two null models. Crosstalks were calculated as described in the main text section “Alteration of intra-cellular crosstalks in cancer active fibroblasts”.

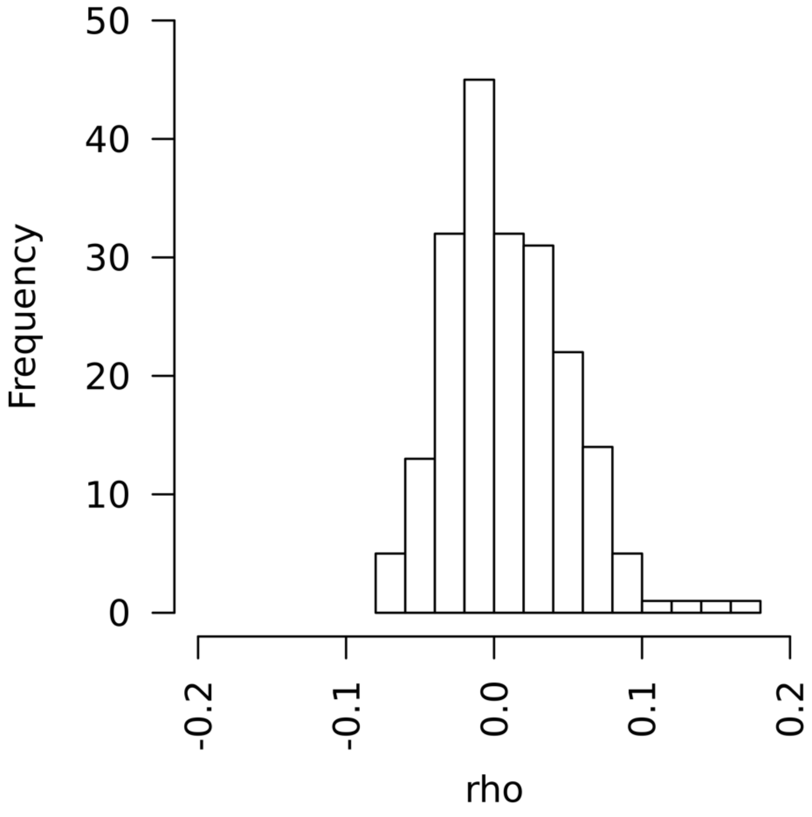

**Supplementary Figure 3**. Comparison of cell clusters and cell type annotation of the 8 TNBC samples used in our proof-of-concept.

On the left, the cell clustering performed by Pal et al. (2021)[3] and provided as the Seurat object "SeuratObject_TNBC.rds"[4]. On the right, the cluster annotation reconstructed by us based on what is written in Pal et al. (2021)[3] and what is available in the Seurat object "SeuratObject_TNBCSub.rds"[4], as described in the main text.

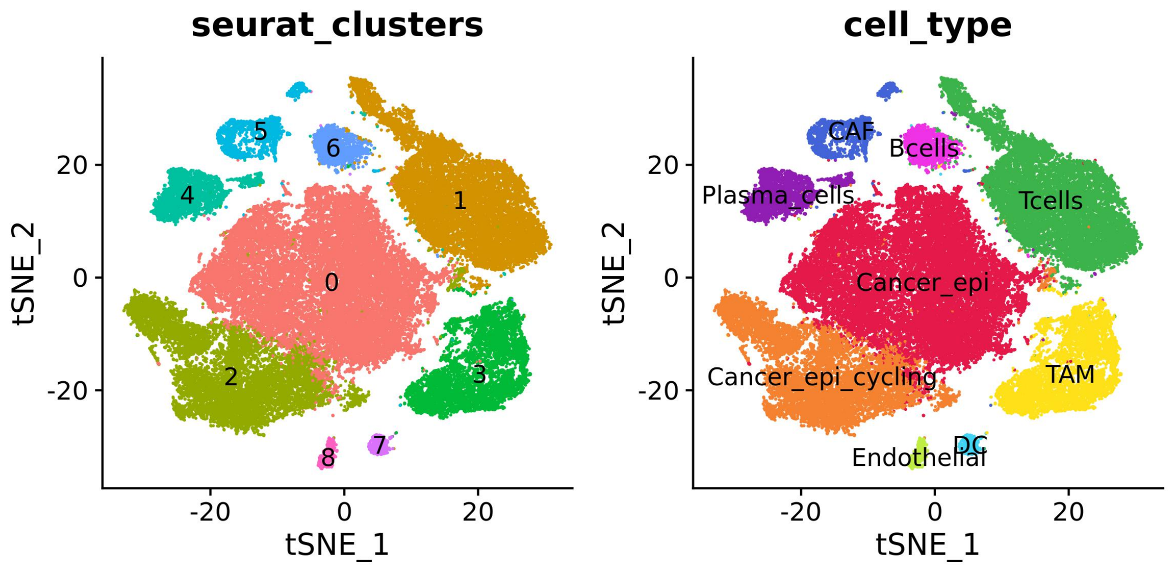

### Supplementary Tables

**Supplementary Table 1**. Comparison between Ulisse and existing approaches for intra-cellular and extra-cellular crosstalks.

As stated in the main text, there are several differences between Ulisse and other existing approaches that focus, in the field of network medicine, on intra-cellular and extra-cellular crosstalks. Many tools focus on cell-cell communication, some of which also consider intra-cellular signalling. Conversely, the availability of computational tools to study pathway crosstalks is limited. Overall, there is a high variability among approaches in terms of focus, mathematics, interaction source, statistical assessment and integration between intra-cellulat and inter-cellular signalling[5]. This variability, as well as the absence of a ground truth, makes the direct comparisons limited and of difficult interpretation[6]. The tools listed in the table are implemented as R packages, apart from CellPhoneDB[7] that is written in python. LR: ligand-receptor; TF: transcription factor.

| ***Approach*** | **Focus** | **Interactions** | **Statistical assessment** | **Integrated crosstalk** |
|---|---|---|---|---|
| *Ulisse* | • Cell clusters<br>• Gene sets<br>• Genes | • Ominpath<br>• STRING<br>• User-provided | • Randomization of gene-gene interactions<br>• Randomization of gene weights | • Crosstalk between genes involved in inter-cellular communications and intra-cellular processes |
| *CellPhoneDB*[7] | • Genes | • built-in<br>• user-provided (intercellular only) | • Permutation of cell labels | • Based on receptor-TF interactions |
| *CellChat*[8] | • Cell clusters<br>• Genes | • Built-in<br>• User-provided | • Based on modelling of LR interactions<br>• permutation of cell labels (cell-cell communication analysis) | • No |
| *SigleCellSignalR*[9] | • Genes | • Built-in | • No | • Expressed genes involved in pathway containing receptors |
| *NicheNet*[10] | • Genes | • Built-in | • No | • LR-transcriptional Regulators-Target genes |
| *PathNet*[11] | • Pathways | • Built-in | • Randomization of gene weights | • No |

**Supplementary Table 2**. Estimated probabilities of the crosstalk values reported in Supplementary Figure S1.

| $X$ | $Y$ | $\rho_{\mathrm{A}}$ | $\rho_{\mathrm{u}}$ | $p$ |
|---|---|---|---|---|
| OXIDATIVE_PHOSPHORYLATION | P53_PATHWAY | 0.799 | 0.108 | 0.298 |
| E2F_TARGETS | G2M_CHECKPOINT | 0.030 | 0.228 | 0.041 |
| MYC_TARGETS_V1 | OXIDATIVE_PHOSPHORYLATION | 0.132 | 0.001 | 0.001 |
| ALLOGRAFT_REJECTION | MYC_TARGETS_V1 | 0.001 | 0.001 | $\sim 10^{-5}$ |

**Supplementary Table 3.** Differential expression analysis results. The table contains the results obtained with "FindAllMarkers()" function form Seurat, together with the cell type annotation and score u used in Ulisse analysis, calculated as described in Method. See file "SupplementaryTables_3-11.xlsx".

**Supplementary Table 4.** Differential expression genes (DEG) mapping data. The table contains data about the number of genes tested and DEGs mapped to STRING, Omnipath networks and Hallmarks gene sets, for each cell type. See file "SupplementaryTables_3-11.xlsx".

**Supplementary Table 5.** Intracellular crosstalk results in Cancer cells. The table contains the analysis results obtained from Ulisse, listing: names of the gene sets ("S1_name", "S2_name"), crosstalk score c, sizes of the gene sets ("S1_size", "S2_size") and the subset involved in the crosstalk calculation ("S1_S2_size", "S2_S1_size" = number of genes; "S1", "S2": gene names; "u1", "u2" = sum of weights u of the genes involved), number of links between the pairs ("dL" = affected; "L" = possible) and their crosstalk saturation ("r_c"), summary score s, and p-values ("pA", "pU" and their combination "p"). The last column ("significance") indicates whether each pair is supported by one ("M_A | M_U < 0.01"), both ("M_A & M_U < 0.01") or no ("M_A & M_U > 0.01") null models. See file "SupplementaryTables_3-11.xlsx".

**Supplementary Table 6.** Over-Representation Analysis (ORA) on cancer cells DEGs results. The table contains the results of the ORA on Hallmarks gene sets, by listing the number of tested genes ("N"), DEGs ("wb"), non-DEG genes ("bb"), genes in the gene set ("bd") and DEGs in gene set ("wbd"), together with expected value ("exp"), enrichment ("er") and statistical evaluation ("p_val", "p_adj", "q_val"). See file "SupplementaryTables_3-11.xlsx".

**Supplementary Table 7.** Gene classification of cancer intracellular crosstalk. The table shows the results of gene classification analysis performed on significant crosstalk, as described in Methods. For each gene there are: crosstalk diversity, saturation, gene sets involved, and gene set measures used for normalization (“ddX”, “rX”, “ddX_S”, “dX”, “dX_S”, respectively); interactor diversity, saturation, genes involved and genes measures used for normalization (“ddA”, “rA”, “ddA_gene”, “dA”, “dA_gene”, respectively); initial weight of the gene (“u”). See file “SupplementaryTables_3-11.xlsx”.

**Supplementary Table 8.** Intercellular crosstalk results. The table contains the results of intracellular crosstalks analysis performed as described in Methods. The data listed are the same in Supplementary Table 5. See file “SupplementaryTables_3-11.xlsx”.

**Supplementary Table 9.** Cancer-CAF intercellular crosstalk details. This table contains data about the communication between cancer cells and CAF, with the detail of the gene-gene and score (“u12”, calculated as product of gene weights) of their interaction. See file “SupplementaryTables_3-11.xlsx”.

**Supplementary Table 10.** Intercellular crosstalk gene classification. Here are listed the results of the gene classification analysis performed on significant intercellular communications, as described in Methods. The data listed correspond to Supplementary Table 7. The last column (“specific”) indicates if a gene is present among cancer cell (“Cancer_epi”), CAF or both communicating gene sets, or none of them (“other”). See file “SupplementaryTables_3-11.xlsx”.

**Supplementary Table 11.** Integrated crosstalk analysis in cancer cells, involved in communication with CAF. The table shows the result of integrated crosstalk analysis. Here, “ccc” columns indicate which communication is analysed, and “S1_name” represent cancer cells genes involved in the communication with CAFs. Data listed in the other columns correspond in meaning to Supplementary Table 5. The last column (“intracellular”) indicates if the pathway in S2_name is involved in significant intracellular crosstalk. See file “SupplementaryTables_3-11.xlsx”.
